# Metagenomic characterization of soil microbial communities in the Luquillo experimental forest (Puerto Rico) and implications for nitrogen cycling

**DOI:** 10.1101/2020.06.15.153866

**Authors:** Smruthi Karthikeyan, Luis H. Orellana, Eric R. Johnston, Janet K. Hatt, Frank E. Löffler, Héctor L. Ayala-del-Río, Grizelle González, Konstantinos T Konstantinidis

## Abstract

The phylogenetic and functional diversity of microbial communities in tropical rainforests, and how these differ from temperate communities remain poorly described but are directly related to the increased fluxes of greenhouse gases such as nitrous oxide (N_2_O) from the tropics. Towards closing these knowledge gaps, we analyzed replicated shotgun metagenomes representing distinct life zones from four locations in the Luquillo Experimental Forest (LEF), Puerto Rico. These soils had a distinct microbial community composition and lower species diversity when compared to temperate grasslands or agricultural soils. Unlike temperate soils, LEF soils showed little stratification with depth in the first 0-30cm, with ~45% of community composition differences explained solely by location. The relative abundances and nucleotide sequences of N_2_O reductases (*nosZ*) were highly similar between tropical forest and temperate soils. However, respiratory NO reductase (*norB*) was 2-fold more abundant in the tropical soils, which might be relatable to their greater N_2_O emissions. Nitrogen fixation (*nifH*) also showed higher relative abundance in rainforest compared to temperate soils (20% vs. 0.1-0.3% of bacterial genomes in each soil type harbored the gene, respectively). Collectively, these results advance our understanding of spatial diversity and metabolic repertoire of tropical rainforest soil communities, and should facilitate future ecological modeling efforts.

**Importance:** Tropical rainforests are the largest terrestrial sinks of atmospheric CO_2_ and the largest natural source of N_2_O emissions, two critical greenhouse gases for the climate. The microbial communities of rainforest soils that directly or indirectly, through affecting plant growth, contribute to these fluxes remain poorly described by cultured-independent methods. To close this knowledge gap, the present study applied shotgun metagenomics to samples selected from 3 distinct life zones within the Puerto Rico rainforest. The results advance our understanding of microbial community diversity in rainforest soils and should facilitate future studies of natural or manipulated perturbations of these critical ecosystems.

## INTRODUCTION

Soil microbiomes are one of the most complex ecosystems owing to microenvironments and steep physicochemical gradients, which can change on a micrometer or millimeter scale (1–3). Temporal and spatial heterogeneity, demographic stochasticity, ecotype mixing, dispersion and biotic interactions are the major drivers of soil microbial diversity in these ecosystems (4, 5). The formation of such “metacommunities” coupled with biogeography and other edaphic factors greatly influence the functional and taxonomic profile of a soil ecosystem at any given location (6).

Tropical rainforests (“forests” hereafter) are characterized by humid and wet climate patterns and account for a large portion of the world’s total forest cover (7). These forests have high levels of primary productivity (~30% of the total global production) due to large amounts of precipitation coupled with year-long warm temperatures and high levels of light (8). Consequently, high levels of biodiversity are observed in these forest soils with unique microbial genotypic signatures being exclusive to this habitat/location, along with only a few cosmopolitan taxa that are shared with other (non-tropical forest) habitats (9, 10). Although tropical forest soils are critical ecosystems that host a plethora of distinct ecological niches, little is known about the metabolic potential of tropical soils, especially, across elevation and depth gradients. Describing this metabolic diversity is important for studying and monitoring the microbial activities related to greenhouse gas fluxes, namely, nitrous oxide (N_2_O) and carbon dioxide (CO_2_) from the tropical soils (11).

Notably, tropical forests represent the largest terrestrial sinks of atmospheric CO_2_ and the largest natural source of N_2_O emissions (12–15). Natural soils have been reported to contribute over 43% of the total global N_2_O emissions, with tropical ecosystems being the highest contributors, having 2 to 4 times higher contributions compared to natural temperate ecosystems (16–19). These soils are also responsible for about 70% of terrestrial nitrogen fixation, which underlies, at least in part, their high rates of net primary productivity (11, 20).

Microbially-mediated nitrification and denitrification are the biotic processes contributing the most to global N_2_O soil emissions (60-70%) (19, 21, 22), although chemodenitrification, i.e., ferrous iron generated by ferric iron-reducing bacteria reacting with nitrite to produce N_2_O abiotically, is also likely high in iron-rich tropical soils (23). In soils, N_2_O is biologically produced as a result of incomplete nitrification, DNRA (dissimilatory nitrite reduction to ammonium) or denitrification respiratory pathways (22, 24, 25). Respiratory nitric oxide reductase (*nor*) is a key contributor to the microbial production of N_2_O and is commonly encoded in the genome of denitrifying bacteria as well as some ammonia-oxidizing organisms (22, 26–30).

While both biotic and abiotic processes contribute to N_2_O production, consumption of N_2_O is exclusively mediated by microbial N_2_O reductase (NosZ) activity (31–34). Yet, whether the denitrifying microorganisms in these soils differ from their counterparts in temperate soils and, if their functional genes present in the community reflect the high nitrogen fluxes, remain unanswered questions despite their apparent importance for better management and modeling of tropical soil ecosystems. It has also been demonstrated that tropical forests have significantly higher rates of nitrogen fixation (~70% of total terrestrial nitrogen fixation) compared to other ecosystems, significantly affecting the nitrogen budgets in these ecosystems (3, 35–37). For instance, higher rates of nitrogen fixation in soils have been linked to nitrous oxide emissions (N loss) due to reduced N retention capacities (11, 38, 39). How these ecosystem rates translate to the nitrogen-fixing microbial (sub)community diversity and gene potential remains unclear.

The Luquillo Experimental Forest (LEF), also known as the El Yunque National Forest in Puerto Rico (PR), has been a long term ecological research (LTER) site since 1988. The site is dedicated to the assessment of the effects of climate drivers on the biota and biogeochemistry. The forest has been subjected to several disturbance regimes over the last few decades, mostly natural and -to a smaller extent-anthropogenic such as tourism and experimental manipulations (40, 41). This site encompasses distinct “life zones” characterized by sharp environmental gradients even across small spatial scales (40, 42, 43). The broad life zones based on the Holdridge classification system include the rain forest, wet forest, lower montane wet forest, and lower montane rain forest. These life zones are distinguished by elevation, temperature and rainfall patterns in addition to other edaphic factors (44–47). The elevation and rainfall patterns also tend to influence oxygen availability, redox potential, nutrient uptake and organic decomposition rates (44, 47, 48). The dynamic interplay of existing physicochemical gradients and climatic factors gives rise to a complex mosaic of biodiversity patterns observed in this forest (45). Hence, LEF represents an ideal environment to study tropical microbial community diversity patterns and their impacts on carbon and nitrogen cycling. The four sampling sites of this study were chosen to represent the distinct vegetation and life zones within the LEF.

Previous studies in the LEF, and similar forest regions, have mostly focused on the effects of redox dynamics, litter decomposition, nitrogen (N) and other nutrient fertilization on microbial community activity through enzyme assays. Few studies have examined microbial diversity patterns across an elevation gradient and were only based on low-resolution techniques such as terminal restriction fragment length polymorphism analysis (14, 49–53). Furthermore, studies linking marker-gene abundances (related to nitrogen cycling) with *in-situ* flux measurements showed very high N_2_O fluxes in the forest soils (54). However, the *nosZ* primers targeted only the typical (Clade I) clades, thereby introducing a primer bias, which can be circumvented by employing metagenomic analyses.

With recent developments in next generation DNA sequencing and associated bioinformatics binning algorithms, near-complete metagenome-assembled genomes (MAGs) can been recovered without cultivation (55, 56), opening new windows into studying soil microbial communities. Here, shotgun metagenomes originating from soils from the four different locations/life zones and three different depths in the LEF were analyzed to describe the microbial community diversity, biogeographical patterns, and metabolic potential differences across samples. Furthermore, the metagenomic data obtained from these soils were also compared to similar data from temperate grasslands in Oklahoma (OK) (57) and agricultural soils from Illinois (IL), USA (56) obtained previously by our team. By analyzing near-complete MAGs, we show that the most abundant microbial population (based on number of reads recruited) at each of the sampling locations represent sequence-discrete populations, similar to those observed in other habitats (58). Using such sequence-discrete populations as the fundamental unit of microbial communities, we subsequently assess the population distribution at high resolution across the sampling sites (biogeography) and the gene content they encoded, with a focus on nitrogen metabolism.

## RESULTS

### Diversity of forest microbial communities

The LEF soil communities were compared to those of intensively studied ecosystems, namely the Oklahoma temperate grassland (OK) (57, 59) and Illinois agricultural soils (IL) (56), which were previously characterized with similar shotgun metagenomics approaches. Shotgun metagenomic sequencing recovered a total of 370 million reads across the 4 sites (Suppl. Table S2). Nonpareil 2.0 (60) was used to estimate sequence coverage, i.e., what fraction of the total extracted community DNA was sequenced. Nonpareil analysis of community diversity (Suppl. Fig. S1) showed that the agricultural Urbana (IL) site had the highest diversity of all the soils compared (NP diversity 24.02; note that NP values are given in log scale), and consequently, the lowest sequence coverage at (only) 37.23%. El Verde and Pico del Este (20-30cm) were the least diverse or most completely sequenced with 87.1% and 73.4% coverage respectively (NP diversity of 19.6 and 20.6 respectively or about 2-3 orders of magnitude less diverse). Overall, OK and IL soils appear to be more diverse than the PR soils by about two orders of magnitude, on average, with an average Nonpareil value of 22.75±0.37. Nearly complete coverage for El Verde and Pico del Este (20-30cm samples) would require 2.402e+09bp and 8.735e+09bp, respectively, and, for the same level of coverage, the more complex communities in Urbana (IL) would require a substantially higher sequencing effort of 1.282e+12bp. The OK soils had an estimated sequencing depth of 2.063e+11± 1.436e+11 bp.

### Community composition variation across the forest sites based on 16S rRNA gene sequences

The number of total 16S-rRNA gene-based OTUs (Operational Taxonomic Unit) observed in each metagenome as well as the Chao1 estimate of total OTUs present reflected the degree of undersampling at each site (Suppl. Fig. S1 and S2), and were also consistent with the Nonpareil coverage estimates (Fig. 1). When Puerto Rico tropical soils (PR) were compared with the agricultural and grassland soils from the United States at the phylum level, *Proteobacteria, Acidobacteria* and *Actinobacteria* were the most abundant taxa across all ecosystems. However, in the forest soils, a few highly abundant OTUs dominated the entire soil community whereas in the OK and IL soils, OTUs were more evenly distributed (Suppl. Fig. S2), consistent with the Nonpareil diversity results. Only 1.28% of the total detected OTUs (out of a total 8019, non-singleton OTUs) were shared among all PR samples, while 49.95% of OTUs were exclusive to a particular sampling site in PR, reflecting partly the under-sampling of the extant diversity by sequencing. Only 0.37% of the OTUs (out of a total 13760, non-singleton OTUs) were shared among all the sites across all 3 ecosystems, all of which were assignable to *Alphaproteobacteria, Acidobacteria, Verrucomicrobia* and *Actinobacteria*.

**Fig. 1:**
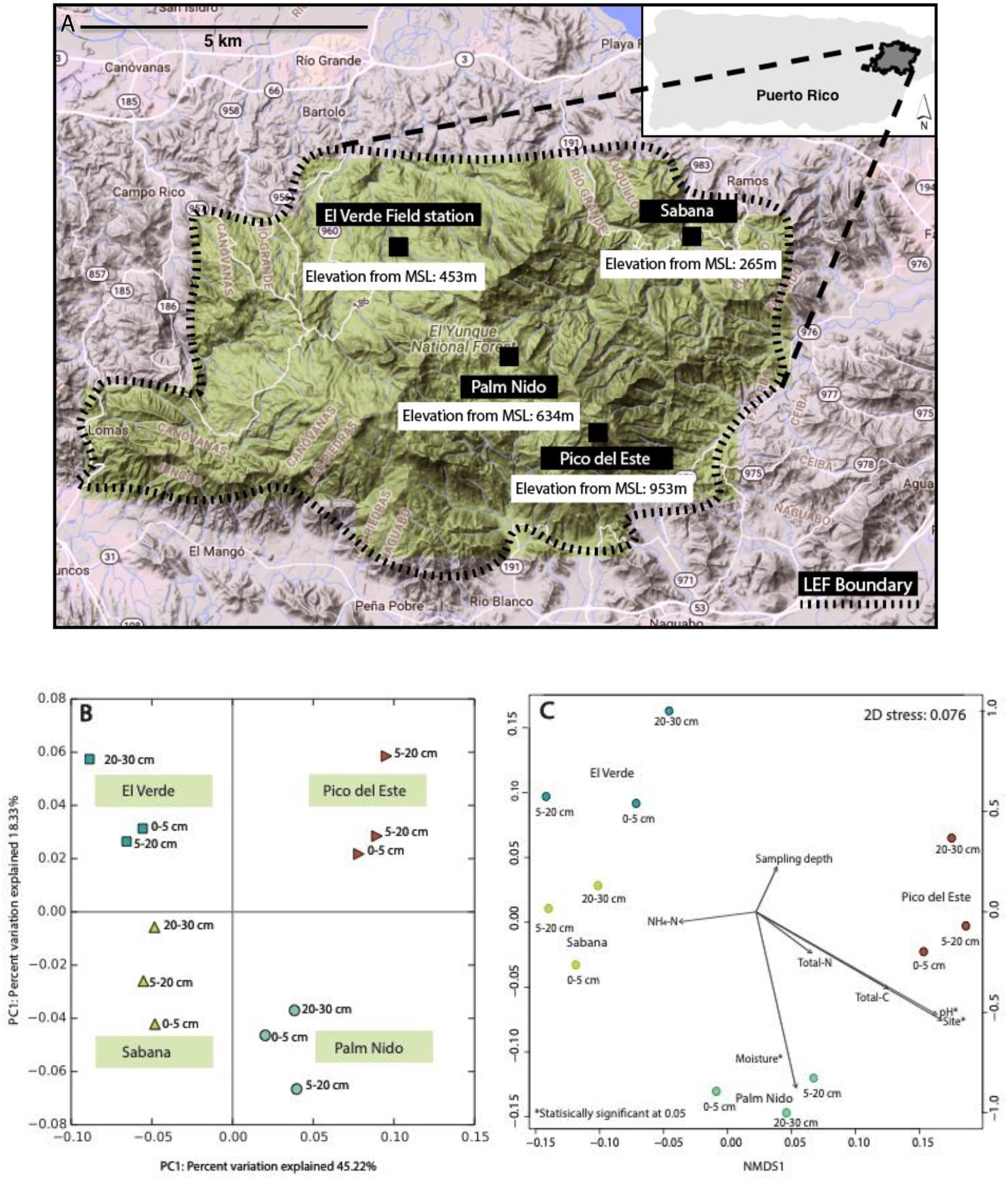
Sampling location map and microbial community diveristy among the study sites. A. Map of the four sampling sites within the Luquillo Experimental Forest (LEF). B. Principal co-ordinate analysis (PCoA) plots based on MASH distances, colored by sampling site, C. Nonmetric multidimensional scaling (NMDS) plot with the soil physicochemical parameters incorporated. The arrow lengths are proportional to the strength of the correlations obtained between measured soil physicochemical parameters and each ordination axis.

Further, applying four additional DNA extraction methods on a selected subset of our samples, including two manual phenol chloroform-based methods that are often advantageous for iron rich soils like those in tropical forest, revealed similar levels of diversity, more or less (Suppl.Fig. S3). Hence, the diversity patterns reported here are robust and independent of the DNA method used.

### Factors driving community diversity in the forest soils: Multidimensional scaling analysis of beta diversity

The PCoA (Principal Coordinate Analysis) plots, constructed based on the MASH distances among whole metagenomes, showed a clustering pattern that was primarily governed by site/location. Accordingly, site explained 45.22% of the total diversity (Fig. 2B). The non-metric multidimensional scaling (NMDS) analysis of the data revealed only site, pH and soil moisture to be statistically significant physicochemical parameters in explaining the observed community diversity (Fig. 2C, Suppl. Table S3). ANOSIM values also indicated site to be a more important factor than depth, with a P value of 0.001 and 0.94, respectively. Based on the distance-based redundancy analysis (dbRDA), site was the most significant factor, even when the interplay between site and sampling depth was accounted for (Suppl. Table S4). Table 1 shows the partitioning of the variance between the proportion that is explained by constrained axes (i.e., environmental variables measured) and the porportion explained by unconstrained axes (i.e., variance not explained by environmental variables measured). The total variance explained by all (measured) environmental variables was 80.2% (Table. 1), which is remarkably high for a soil ecosystem (61).

**Table 1:** Proportion of total microbial community diversity explained by measured soil environmental factors.

|  | Inertia | Proportion | Rank |
| --- | --- | --- | --- |
| Total | 0.1092 | 1 |  |
| Constrained | 0.0876 | 0.8021 | 6 |
| Unconstrained | 0.02161 | 0.1978 | 5 |

Site, sampling depth, pH, total nitrogen, total carbon, moisture data were considered in the analysis

**Fig. 2:**
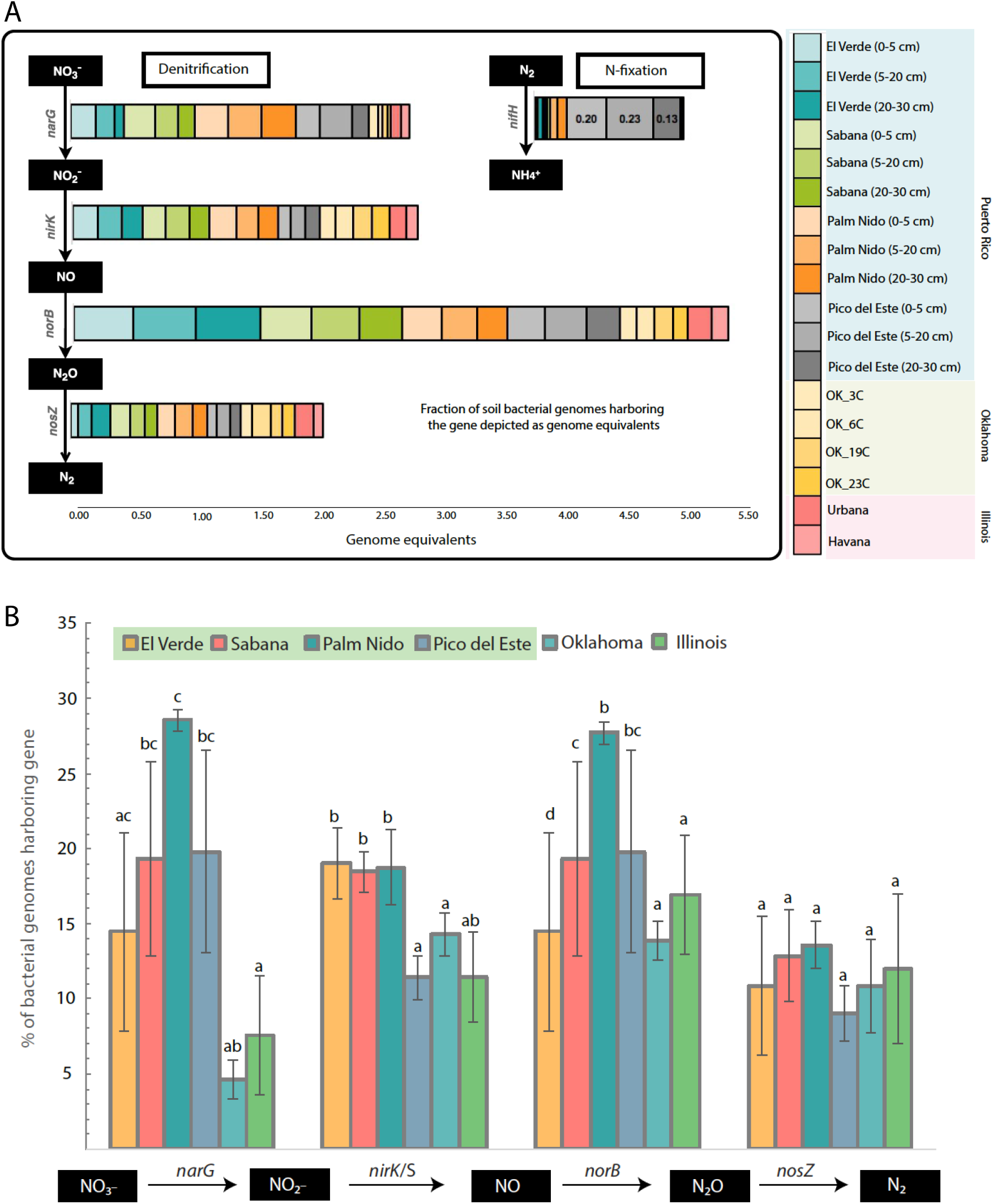
Abundance of N cycling genes and their distribution across soil ecosystems. A. Abundance of hallmark genes for denitrification, DNRA and nitrogen fixation pathways, represented as genome equivalents (% of total bacterial genomes sampled that carry the gene) in the metagenomes studied (see Figure key). B. Frequency of genomes carrying the respective denitrifying gene across the three ecosystems studied. Genes denoted by the same letter are not statistically significantly different between ecosystems (ANOVA Tukey test). Statistical significance reported at p < 0.05. Note that nitrification genes were not detected in any of the Puerto Rico sites.

### Major N cycling pathways

Genes encoding proteins involved in denitrification and nitrogen fixation were the most abundant nitrogen (N) cycling pathway genes detected at different sites. Overall, the forest soils harbored about a 2-3-fold higher abundance of denitrification genes, i.e., *narG, nirK*, and *norB* catalyzing the reduction of nitrate, nitrite, and nitric oxide, respectively, compared to the grassland and agricultural soils (Fig. 2A). For instance, the *norB* gene abundance was found to be at the highest abundance among the denitrification genes, with ~37% (SD 9.5%) of the genomes in the PR soils predicted to contain a *norB* gene, compared to ~17% (SD 4%) and ~14% (SD 1.3%) at IL and OK, respectively. Similarly, *narG* showed a 3-fold higher abundance in the PR soils compared to IL and OK soils (Fig.2B). While denitrification gene abundances appeared higher in the tropical soils, the relative abundance of *nosZ* gene, i.e., 11.6% (SD 3%) of the total genomes across the four locations in the LEF were predicted to encode *nosZ*, similar to *nosZ* relative abundance in IL and OK soils, i.e., 11.75% (SD 5%) and 11.08% (SD 3%), respectively (not statistically significant at p=0.05). Similar to *nosZ*, DNRA gene abundances (namely, *nrfA*) was similar across all sites studied herein (9%, SD 1.9%).

### Predominant NosZ clades are shared among soil ecosystems

Placing *nosZ*-encoding reads to a reference *nosZ* phylogenetic tree revealed that atypical clades (clade II *nosZ)*, affiliated predominantly with *Opitutus, Anaeromyxobacter* and other closely related genera, dominated the *nosZ* gene pool in the tropical forests (Figs. 3, Suppl. Figs.S4-S7). In contrast, a very small fraction of reads (<10% of total *nosZ* reads) were recruited to typical *nosZ* clades (or clade I). Members belonging to the clade II *nosZ* dominated the *nosZ* gene pool in OK and IL soils as well, with IL agricultural soils showing the greatest *nosZ* sequence diversity among the three regions. Notably, *O. terrae*-affiliated sequences represented the most abundant sub-clade (*nosZ* OTUs/sub-clades were defined at the 95% nucleotide sequence identity level) in all regions. Furthermore, most of the *O. terrae*–affiliated reads in the forest soil dataset appeared to be assigned to a single sub-clade, while their counterparts in the OK and IL soils appeared to be more evenly distributed among several closely related *nosZ* sub-clades, i.e., showing higher sequence diversity (Fig. 3, Suppl. Figs. S4-S7)*. O. terrae* (strain DSM 11246/PB90-1) *nosZ* reads at >95% identity made up between 20% and 60% of the total *nosZ* reads recovered from the El Verde site and, together with the second most abundant sub-clade from *Anaeromyxobacter sp.*, contributed over 30% of the total *nosZ* reads across all four PR locations (Fig. 5). Despite the significant taxonomic diversity observed in these soils (Suppl Fig. S2), the soils from PR shared several abundant *nosZ* gene sequences/sub-clades at >95 nucleotide identity with soils in OK and IL (Fig. 3). Furthermore, in order to compare the predominant *nosZ* clades across the samples shown here, a new phylogenetic reference tree was constructed based on almost full length sequences obtained from the assemblies/MAGs obtained from the metagenomes studied here (namely PR,OK,IL). The short-reads identified as *nosZ* from the PR soils were placed on this tree and show that the majority of these reads are recruited by the *nosZ* sequences obtained from these assemblies/MAGs, indicating that the *nosZ* sequences across these ecosystems studies here are similar (Suppl. Fig. S8)

**Fig. 3:**
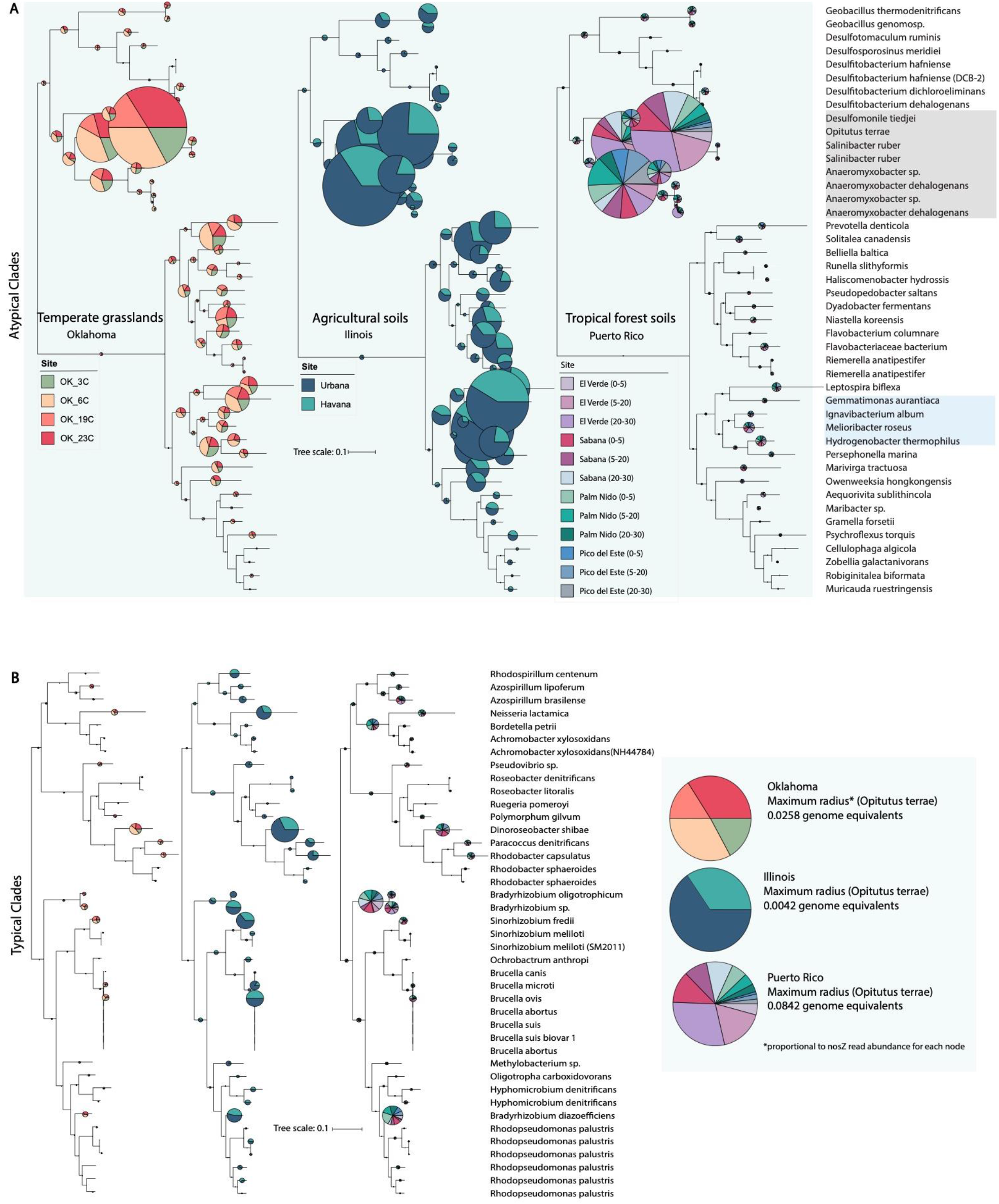
Phylogenetic diversity of *nosZ*-encoding sequences recovered in each soil ecosystem. *nosZ* sequences were identified by the ROCker pipeline and placed in a reference *nosZ* phylogeny as described in the Materials and Methods section. The radii of the pie charts are proportional to the number of reads assigned to each sub-clade and the colors represent the sampling sites from each ecosystem (see Figure key). Sub-clades highlighted in grey indicate the most abundant sub-clades across all three ecosystems whereas the ones highlighted in blue were abundant only in agricultural soils (IL). A. *nosZ* reads from every sampling site recruiting to atypical (Clade II) clades. B. *nosZ* reads recruiting to typical (Clade I) clades. Inset shows the most abundant sub-clade (*Opitutus terrae*) from panel A and its distribution across all sites. Note that in all three ecosystems most of the reads recruit to atypical sub-clades. Suppl. Fig. S7 shows the distribution of the reads among the most abundant sub-clades in detail.

### Nitrogen fixation potential

The nitrogen fixation genes (mainly *nifH*) were present at a much lower abundance in the lower altitude forest samples. For instance, only ~1-3% of all genomes in the lower altitude samples were predicted to encode *nifH* compared to a ~20% of the genomes in the higher elevation samples (Pico del Este) (Fig. 2A), and almost none of the reads from IL and OK metagenomes appeared to encode *nifH* (<0.1%). Therefore, nitrogen fixation gene abundance patterns indicated a much stronger selection for nitrogen fixation in the tropical forest relative to temperate agricultural or natural prairie soils, especially at higher elevations. Furthermore, no ammonia oxidizing genes (*amoA*) were detected in any of the soils except for Urbana soils (IL), which had a history of fertilizer (N) input.

### Recovery of metagenome-assembled genomes (MAGs) representative of each site

In order to test the effect of biogeography (i.e., limits to dispersion) of taxa across the elevation gradient sampled, the distribution of abundant MAGs recovered from each PR sampling site \–(assembly and MAG statistics provided in Suppl. Table S6) were assessed across the sites using read-recruitment plots (62). Taxonomic assignment using the Microbial Genomes Atlas (63) revealed that the most abundant MAG at site El Verde (lowest elevation), representing 4.39% of the total metagenome, and was affiliated with an unclassified *Verrucomicrobia*. The second most abundant (1.8% of total) was likely a member of the genus *Ca. Koribacter* (*Acidobateria*) followed by an unclassified member of *Acidobacteria* (1.45% of total). The *Verrucomicrobium* MAG was found at an abundance of 1.03% of the total population at Sabana, and at 0.07% and 0.03% in Palm Nido and Pico del Este (highest elevation), respectively. Uneven coverage across the length of the reference sequence and nucleotide sequence identities were observed in the recruitment of short-reads from Palm Nido and Pico del Este as well as with all OK datasets, indicating that the related populations in the latter samples were divergent from the reference MAG (Suppl. Fig. S10). Therefore, at least this abundant low-elevation Verrucomicrobial population did not appear to be widespread in the other samples analyzed here (Suppl. Fig. S10). Similarly, the other abundant MAGs from other sites in the forest soils were unique to the corresponding sites (elevation) from which they were recovered. Almost all MAGs used in the analyses were assignable to a novel family, if not higher taxonomic rank, according to MiGA analysis (when compared to 11,566 classified isolate genomes available in the NCBI prokaryotic genome database), underscoring the large unexplored diversity harbored by the PR tropical rainforest soils. The sequence diversity/complexity as well as sequencing depth limited large-scale recovery of high-quality MAGS.

### Functional gene content of the MAGs

The genome sequences of the most abundant MAGs from each location (n=6) were analyzed in more detail to assess the functions they encoded, especially with respect to N cycling pathways (Fig. 4). MAGs from Pico del Este (highest elevation) showed a high abundance of N metabolism related genes compared to MAGs from other sites (Fig. 4). Most notably, genes related to nitrogen fixation were found only in the Pico del Este MAG, which was consistent with the short read analysis datasets showing greater relative abundance of *nifH* at this site. Nitrification (ammonia oxidation related genes) gene clusters were not detected in any of the recovered MAGs. *norB* and *nosZ* genes were found in three out of the six abundant MAGs analyzed. The most abundant El Verde MAG, most closely related to *O. terrae* (AAI = 40 %), possessed a *nosZ* gene, which was congruent with the *nosZ* phylogeny described above (i.e., ~60% of the *nosZ*-encoded reads from El Verde had a closest match to *O. terrae nosZ* sequences).

**Fig. 4:**
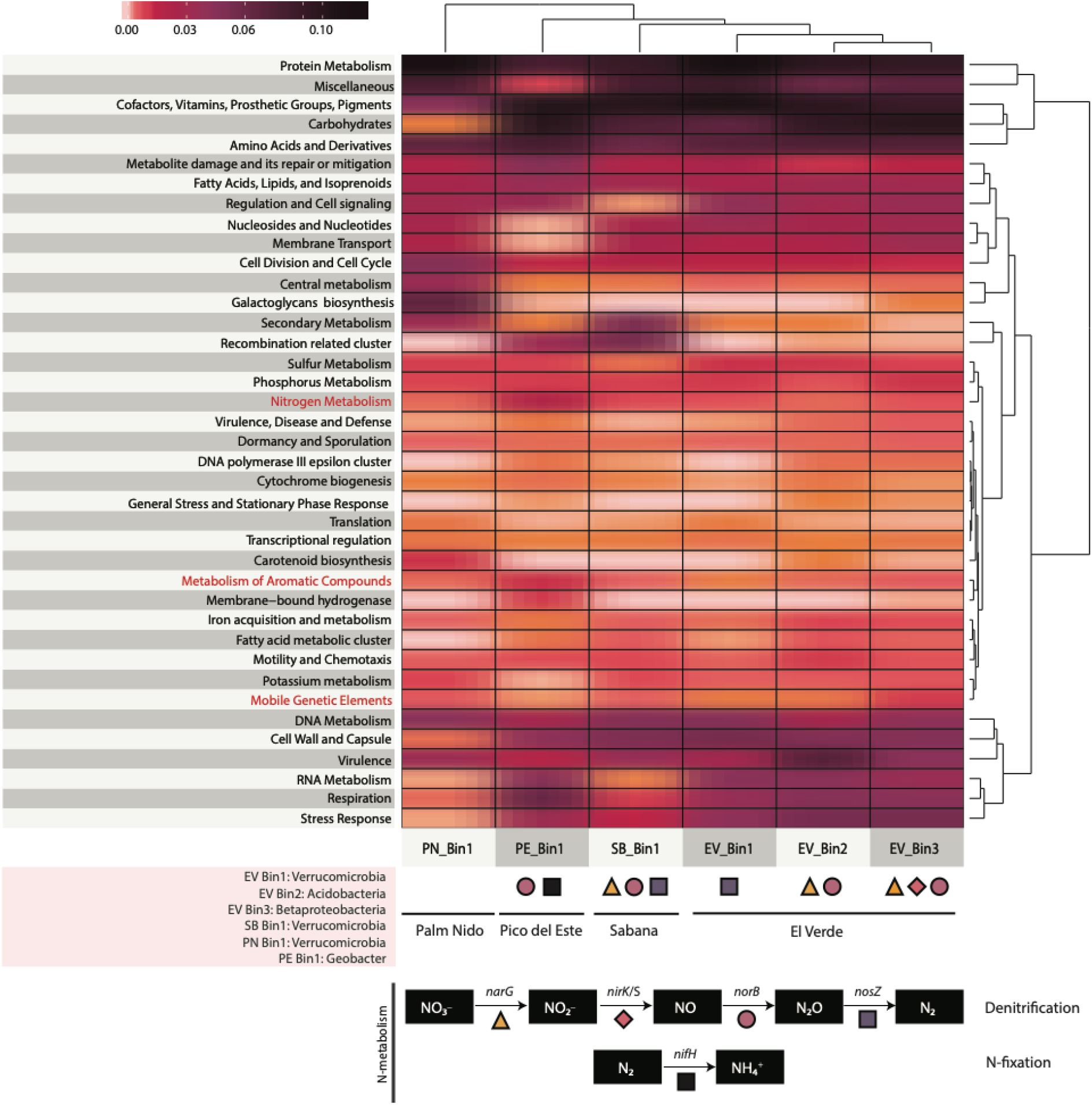
Functions encoded by the recovered population MAGs. Heatmap showing the relative abundance of genes encoding the major metabolic functions (Level 1 of the SEED subsystem category) for each MAG recovered from the four sites in Puerto Rico. The taxonomic classification of each MAG based on MiGA is shown on the bottom left. The symbols at the bottom of the heatmap denote the presence (or absence) of specific N-cycling genes, namely denitrification and nitrogen fixation. No genes involved in nitrification were detected in any of the bins.

## DISCUSSION

The present study reported the taxonomic and gene content diversity of the poorly characterized tropical rainforest soils by using whole-community, shotgun metagenomic sequencing of samples from the El Yunque forest, Puerto Rico. The recovered near-complete MAGs represented several abundant and widespread organisms within this ecosystem that could serve as model organisms for future studies. Furthermore, since the Luquillo Experimental Forest (LEF) within El Yunque is subjected to varying natural as well as experimental (e.g., warming, phosphorus fertilization) perturbations, our study could also provide a baseline for these perturbations and future soil microbial studies at LEF. Our results revealed that the LEF soils harbor distinct microbial communities at sites with distinct elevation from sea-level. In contrast, and unlike several other soil ecosystems, sampling depth did not have a substantial impact on structuring community diversity, revealing no depth stratification in the LEF soils, at least for the depths sampled here (5-30cm). This could be due to the lack of distinct soil horizons within the first 30cm of the sampling sites, and indicates that the soil formation and/or physicochemical properties in these ecosystems could differ markedly from those in their temperate counterparts (44).

A recent study examining the dominant bacterial phylotypes across the globe found that the predominant phylotypes were widespread across ecosystems. The only exception to this pattern was the forest tropical soils which harbor distinct phylotypes (10). Consistent with these conclusions, the MAGs recovered from each LEF site represented at least novel species and genera, further underlining the under-tapped microbial diversity harbored by tropical forest soils. Currently, the environmental factors driving these diversity patterns remain poorly understood for tropical forest soils (10), but our study provided several new insights into this issue.

In particular, sites El Verde and Sabana (lowest elevation sites) had similar community structure and diversity compared to the two higher-elevation sampling sites with certain MAGs being present at both sites but not in any of the other (higher-elevation) sites examined. This is presumably attributable to both sites having similar climate and vegetation patterns (i.e., Tabonuco forest). On the other hand, Pico del Este was the highest elevation site and experiences almost continuous cloud cover as well as horizontal precipitation. The unique topology of Pico del Este was reflected in distinct and deeply novel MAGs and gene content, which differed markedly from the other three sampling sites within the LEF (PCoA plots, Fig. 2B). The high water content of the Pico del Este soils gives rise to a unique ecosystem dominated by epiphytes (e.g., moss) (64). The epiphytic community has presumably significant impacts on nutrient (e.g., nitrogen) cycling (65), and influences the water input to the soil, thereby shaping a unique habitat/niche for the soil microbes. Free-living microbes have been shown to be one of the highest contributors to biological N fixation in these forests with high rates of nitrogenase activity associated with the presence of moss/epiphytes (53, 66). Consistent with these previous results and interpretations, the Pico del Este showed an extremely high potential for nitrogen fixation, i.e., it was estimated that 1/5 of the total bacterial genomes sampled possessed genes for N fixation, which is at least 10 times greater than any other site evaluated herein. Accordingly, we found that site (location) alone explained about half (45%) of the beta diversity differences observed among the four sampling sites, which reached ~80% when a few physicochemical parameters namely pH and moisture were also included in the analyses (Fig. 2B, Table 1). This is a remarkably high fraction of beta diversity explained by measured parameters for a soil ecosystem (61) and likely reflected that location and the physical properties that characterized different locations within LEF structured diversity much stronger than in other soil ecosystems. Tropical forests have also been shown to have significantly higher rates of nitrogen fixation compared to other ecosystems, which can exceed the N retention capacity of the soil resulting in large N loss as N2O (67). The findings reported here on denitrification gene abundances were generally consistent with these previous observations as well.

Links between soil community structure and nitrogen cycling can help close the knowledge gaps on how the forest ecosystems impact the release and mitigation of certain highly potent greenhouse gases such as N_2_O. The gene abundances observed here, e.g., more than two-fold higher abundance of *norB* (associated with NO reduction to N2O) and similar *nosZ* (N2O consumption) abundances in tropical soils relative to temperate soils were consistent with higher N2O emissions observed previously from the tropics. Further, in acidic soils such as the tropical forest soils evaluated in this study, lack of N limitation can suppress complete denitrification, thereby leading to higher N2O release compared to other soil ecosystems (35). These interpretations were consistent with our observation that the PR soils harbored a relatively high abundance of respiratory (related to denitrification) *norB* genes as well. Previous studies have also suggested that most denitrifying bacterial genomes possess the genes required to reduce nitrate to nitrous oxide but do not possess the gene responsible for the last step i.e., N_2_O reduction to N_2_, leading to the release of N_2_O gas (Braker and Tiedje, 2003; Richardson et al., 2009; Giles et al., 2012; (22, 26–29), consistent with the findings of our study.

It has been established that tropical forest soils are the single highest contributor of natural N2O emissions. While several abiotic and microbial processes can contribute to soil N2O, N2O consumption is an exclusively microbial process, catalyzed by the enzyme product of the *nosZ* genes (34). Based on the assessment of the *nosZ* gene phylogeny, it appears that almost all of the *nosZ* genes from the tropical forest soils studied here belong to a previously overlooked Clade II or atypical *nosZ* genes (32, 34, 68). This clade consists mainly of non-denitrifying, and secondary denitrifying N_2_O reducers. Despite the unique phylogenetic diversity harbored by tropical soils in general, the *nosZ* gene sequence diversity appears to be shared between temperate and agricultural soils (Fig 4). These findings imply strong selection pressure for conservation of nitrous oxide reductase sequences across tropical and temperate soil ecosystems that are not apparently applicable to other N-cycling genes and pathways, which warrants further attention in the future.

Integration of functional (e.g., gene expression) data with *in-situ* rate measurements will provide a more complete picture of the composition and functioning in tropical forest soils. The identification of certain biomarker genes such as *nosZ* sequences in our study could facilitate future investigations on biogeochemical N-cycling and greenhouse gas emissions. For instance, the assembled MAGs and gene sequences provided here could be useful for the design of specific PCR assays for assessing transcript levels (activity), allowing potential linking of carbon dioxide, methane, nitrogen, SOM, etc. turnover to the activity of individual populations. It would also be interesting to assess how the findings reported here for the LEF apply (or not) to other tropical forests especially because our study is based on a relative small sample size. While the diversity in the Puerto Rico soils appears to be lower than that in temperate grassland and agricultural soils, and different DNA extraction methods, including phenol-chloroform- and kit-based, provided for similar results (Fig. S3), it is important to note that DNA of the temperate soil samples was extracted using different methods (OK soils were extracted using the PowerSoil kit). Therefore, it would be important to confirm these preliminary findings by using the exact same DNA extraction and sequencing procedures in all soils. Despite the sample size, however, our results showed differences along the elevation gradient sampled at the LEF that are independent of DNA extraction (Suppl.Fig. S3) or sequencing methods, and consistent with our metadata (Fig. 2), and previous process rate measurements. As the gradients at the LEF also provide a natural setting to interpret the potential ramifications of climate change scenarios such as altered participation patterns, the DNA sequences provided here could facilitate future manipulation experiments with an emphasis on understanding and predicting the effects of climate change on microbial community dynamics along the elevation gradient.

## MATERIALS AND METHODS

### Sampling sites

Soil samples were collected on February 2016 from four locations/sites across the LEF (18.3’ N, 65.80’ W). The four sites namely, Sabana, El Verde field station, Palm Nido and Pico del Este, each located at different elevations from the mean sea level, i.e., 265, 434, 634 and 953 m, respectively, were chosen due to their unique landscape and rainfall patterns, thereby creating distinct ecological niches (Fig. 2A).

The El Yunque forest is categorized into four distinct vegetation zones namely, the Tabonuco, Palo Colorado, Sierra Palm and Dwarf/Elfin forests. Site Sabana and El Verde, which are located at the lowest elevation among the four sites within the LEF, fall under the Tabonuco forest category in terms of vegetation, dominated by the tree species *Dacryodes excelsa* (native to Puerto Rico). They are characterized by canopy cover and low light intensities at the ground level which account for the sparsely vegetated forest floor. However, these sites still harbor the richest flora of all sites (69). Palm Nido is characterized by unstable, wetter soils, steeper slopes and the vegetation is dominated by the Sierra Palm (*Prestoea montana*). The site at the highest elevation, Pico del Este (dwarf forest ecosystem or “elfin woodlands”) is characterized by higher winds, lower temperatures and the vegetation is enveloped by clouds (41, 70) and its main vegetation is comprised of moss and epiphytes. Furthermore, highly acidic soil and continuously water-saturated soils deficient in oxygen are some major characteristics of this ecosystem with most mineral inputs for plants become dissolved in the rain and fog.

Three adjacent soil profiles were taken from each of the four LEF sites (4 sites encompassing 3 lifezones, Palo Colorado was not sampled). For each profile, individual soil cores were taken at each depth (0-5cm, 5-20cm, 20-30cm) using a 3-cm diameter x 15-cm length soil corer (AMS Inc, Idaho) that was decontaminated between samplings by washing with 70% ethanol. Soil samples were stored in sterile Whirl-pak bags and kept on ice during transport and until storage at −80° C. The three cores at each sampling depth were pooled together for community DNA extraction, producing a total of twelve samples across the four sites.

Soil pH was determined using an automated LabFit AS-3000 pH Analyzer, and soil extractable P, K, Ca, Mg, Mn, and Zn were extracted using the Mehlich-1 method and measured using an inductively coupled plasma spectrograph at the University of Georgia Agricultural and Environmental Services Laboratories (Athens, GA, USA). Soil extractable P using this method is interpreted as the bioavailable fraction of P. NH_4_-N and NO_3_-N were measured by first extracting them from soil samples with 0.1 N KCl, followed by the colorimetric phenate method for NH_4_ ^+^ and the cadmium reduction method NO_3_. The physicochemical conditions at the sites during the time of sampling are provided in Supplementary Table (S1).

### Community DNA extraction and sequencing

Total DNA from soil was extracted using the FastDNA SPIN KIT (MP Biomedicals, Solon, OH) following manufacturer’s procedure with the following modifications (71). Soils were air dried under aseptic conditions followed by grinding employing a mortar and pestle. Cells were lysed by bead beating and DNA was eluted in 50 μl of sterile H_2_O. DNA sequencing libraries were prepared using the Illumina Nextera XT DNA library prep kit according to manufacturer’s instructions except the protocol was terminated after isolation of cleaned double stranded libraries. Library concentrations were determined by fluorescent quantification using a Qubit HS DNA kit and Qubit 2.0 fluorometer (ThermoFisher Scientific), and samples were run on a High Sensitivity DNA chip using the Bioanalyzer 2100 instrument (Agilent) to determine library insert sizes. An equimolar pool of the sequencing libraries was sequenced on an Illumina HiSeq 2500 instrument (located in the School of Biological Sciences, Georgia Institute of Technology) using the HiSeq Rapid PE Cluster Kit v2 and HiSeq Rapid SBS Kit v2 (Illumina) for 300 cycles (2 x 150 bp paired end). Adapter trimming and demultiplexing of sequenced samples was carried out by the HiSeq instrument. In total, 12 metagenomic datasets were generated (3 per site for the three depths), and statistic details on each dataset are provided in Supplementary Table S2.

In order to test for any DNA extraction biases of the kit used above, especially for the high iron/clay content that characterizes tropical forest soils and is known to affect the extraction step, four additional DNA extraction methods were performed in parallel on a small subset of samples collected in 2018 from the same sites (6 samples per extraction method for 5 ecxtraction methods covering the 4 sites). The methods included two manual (as opposed to kit-based) phenol-chloroform based methods (72, 73) as well as two other kit-based methods namely; DNeasy PowerSoil and DNeasy PowerSoil Pro (Qiagen Inc.). For this evaluation, the soils were first homogenized and subsequently in five subsamples to use with each method (including the FastDNA SPIN KIT-based method mentioned above). The libraries were constructed and sequenced the same way as described above for the FastDNA SPIN KIT method.

All metagenomic datasets were deposited in the European Nucleotide Archive (ENA) under project PRJEB26500. Additional data is available at http://enve-omics.ce.gatech.edu/data/prsoils.

### Bioinformatics analysis of metagenomic reads and MAGs

The paired end reads were trimmed and quality checked using the SolexaQA (74) package with a cutoff of Q>20 (≥99% accuracy per base-position) and a minimum trimmed length of 50 bp.

#### i) Assembly and population genome binning

Co-assembly of the short reads from the same location was performed using IDBA-UD (75) and only resulting contigs longer than 500 bp in length were used for downstream analysis (e.g. functional annotation and MyTaxa classification). Genes were predicted on the co-assembled contigs using MetaGeneMark (76) and the predicted protein-coding regions were searched against the NCBI All Genome database using Blastp (77). Since the assembly of individual datasets resulted mostly in short contigs (data not shown), the contigs from the co-assembly (combining metagenomes from the three sampling depths, for each site) were used for population genome binning. Contigs longer than 1Kbp were binned using MaxBin (78) to recover individual MAGs (default settings). The resulting bins were quality checked for contamination and completeness using CheckM (79), and were further evaluated for their intra-population diversity and sequence discreteness using fragment recruitment analysis scripts as part of the Enveomics collection (62) essentially as previously described (80).

#### ii) Functional annotation of MAGs

Genes were predicted for each MAG using MetaGeneMark and the predicted protein-coding regions were searched against the curated Swiss-Prot (81) protein database using Blastp (77). Matches with a bitscore higher than 60 or amino acid identity higher than 40% were used in subsequent analysis. The Swiss-Prot database identifiers were mapped to their corresponding metabolic function based on the hierarchical classification subsystems of the SEED subsystem category (Level 1) (82). The relative abundance of genes mapping to each function was calculated based on the number of predicted genes from each MAG assigned to the function (for read-based assessment, see below). Relative abundance data were plotted in R using the “superheat” package (https://arxiv.org/abs/1512.01524). Individual biomarker genes for each step of the nitrogen cycling pathway were manually verified by visually checking the alignment of the identified sequences by the pipeline outlined above against verified reference sequences.

#### iii) Functional annotation of short reads

Protein-coding sequences present in short reads were predicted using FragGeneScan (83) using the 1% Illumina error model. The predicted genes were then searched against the Swiss-Prot database using Blastp (best match). Low quality matches (bitscore < 60) were excluded, and relative abundance of genes mapping to each function was determined as described in the previous section.

### Community diversity estimation

#### i) Nonpareil

Nonpareil (60) was used to estimate sequence coverage, i.e., what fraction of the total extracted community DNA was sequenced and predict the sequencing effort required to achieve “nearly complete coverage”(≥95%). The default parameters in Nonpareil were used for all datasets. Only one of the two paired reads (forward) for each dataset was used to avoid dependency of the paired reads, which can bias Nonpareil estimates (60).

#### ii) MASH and multidimensional scaling

MASH, a tool employing the MinHash dimensionality reduction technique to compare sample-to-sample sequence composition based on k-mers (84), was used to compute pairwise distances between whole metagenomic datasets and construct the distance matrix to be used in multidimensional scaling. Pairwise MASH distances between the metagenomic datasets were computed from the size-reduced sketches (default parameters). PCoA (Principal coordinate analysis) and NMDS (Non-metric multidimensional scaling) were employed to visualize the distance matrix and evaluate the physicochemical parameters driving community diversity, respectively. Furthermore, dbRDA (distance based redundancy analysis), was used to obtain a finer resolution on the observed compositional variation. All of the above startistical analysis were performed using the vegan package in R (85), with default settings.

#### iii) 16S rRNA gene fragments recovered from shotgun metagenomes

16S ribosomal rRNA (16S) gene fragments were extracted from the metagenomic datasets using Parallel-META (86). 16S-carrying reads were classified taxonomically using the GreenGenes database.

Recovered 16S fragments were clustered (‘closed-reference OTU picking’ strategy using UCLUST (87)) and taxonomically classified based on their best match in the GreenGenes database (88) at an ID ≥ 97% in QIIME (89, 90). The relative abundance of the OTUs were calculated based on the number of reads assigned to each OTU. Community composition was assessed based on OTU taxonomic assignments at the genus and the phylum ranks and was compared between the sites based on the relative abundance of OTUs at each site.

### Identification of N cycling genes using ROCker

ROCker (91) was employed for a precise identification and quantification of *nosZ* (encoding nitrous oxide reductase), *norB* (encoding respiratory nitric oxide reductase, cytochrome bc complex associated), *nirK* (encoding nitrite reductase), *narG* (encoding nitrate reductase), *nrfA* (encoding nitrite reductase, DNRA related) *amoA* (encoding ammonia monooxygenase) and *nifH* (encoding nitrogenase) encoding metagenomic reads (http://enve-omics.ce.gatech.edu/rocker/models). Briefly, the short-read nucleotide sequences were searched (using Blastx) against a training set for each abovementioned protein; training sets were manually curated to encompass experimentally verified reference sequences as suggested previously (91). The resulting matching sequences were then filtered using the ROCker compiled model (model for 150bp-long reads for PR and OK soils and 100 bp model for IL soils). Protein abundances (based on the number of reads assigned to the protein) were normalized by calculating genome equivalents. For the latter, the ROCker-filtered read counts were normalized by the median length of the sequences of each protein reference, and the corresponding genome equivalents were calculated as the ratio of NosZ (or another protein of interest) read counts to the RNA polymerase subunit B (*rpoB*), a universal single copy marker, read counts.

### NosZ phylogenetic analysis

The NosZ reference protein sequences were aligned were aligned using CLUSTAL Omega (92) and a maximum likelihood reference tree was created using RAxML v 8.0.19 (93) with a general time reversible model option, gamma parameter optimization and ‘-f a’ algorithm. The ROCker identified NosZ-encoding reads were extracted from all datasets, translated into protein sequences using FragGeneScan, and then added to the reference alignment using Mafft (94). The reads were placed in the phylogenetic tree using RAxML EPA algorithm and visualized using iTOL (95).

### Intra-population diversity assessment based on recovered MAGs

The taxonomic affiliation of individual contig sequences of a MAG was evaluated based on MyTaxa, a homology based classification tool (96). The MiGA (Microbial Genomes Atlas, www.microbial-genomes.org) webserver was used for the taxonomic classification of the whole MAG using the ANI/AAI concept. To assess intra-population diversity and sequence discreteness, each target population MAG was searched against all the reads from each location by Blastn (only contigs longer than 2Kbp were used). Fragment recruitment plots were constructed based on the Blastn matches (threshold values: nucleotide identity ≥75% and alignment length ≥80bp) using the Enveomics collection of scripts (62). The evenness of coverage and sequence diversity of the reads across the length of the reference genome sequence were used to evaluate the presence and discreteness of the population in the chosen dataset.

## Acknowledgments

This work was supported by the U.S. Department of Energy, Office of Biological and Environmental Research, Genomic Science Program (award DE-SC0006662) and US National Science Foundation (award 1831582). GG was supported by the Luquillo Critical Zone Observatory (National Science Foundation grant EAR-1331841) and the Luquillo Long-Term Ecological Research Site (National Science Foundation grant DEB-1239764). All research at the USDA Forest Service International Institute of Tropical Forestry is done in collaboration with the University of Puerto Rico. We thank María Rivera and Humberto Robles from IITF for their help in soil sampling.

